# Sequence-dependent modulation of hepatorenal biochemical markers following Artemether–Lumefantrine and Sulfadoxine–Pyrimethamine exposure in Wistar rats

**DOI:** 10.64898/2026.06.16.730498

**Authors:** Inimfon A. Udoubom, Oboso E. Etim, Jessie I. Ndem, Anietie A. Akpan, Grace E Agu, Ubokutomabasi I. Jonah, Ekpono-Abasi U. James, Ineza Patrick

## Abstract

Malaria remains one of the most pressing health problems, especially in Sub-Saharan Africa. Various antimalarial drugs, used to combat this debilitating illness, may directly or indirectly affect blood indices in humans. This study aims to evaluate the toxicological effects of sequential administration of Artemether-lumefantrine and sulfadoxine-pyrimethamine in male Wistar rats. Thirty (30) mature male Albino Wistar rats weighing between 190-280g were randomly divided into five groups comprising six (6) rats each. Group 1 served as control, Group 2 received Artemether-lumefantrine (8 mg/kg/bw) for 3 days, Group 3 received sulfadoxine-pyrimethamine (0.079 mg/kg/bw) for 1 day, Group 4 received a sequential dose of Artemether-Lumefantrine for 3days and sulfadoxine-pyrimethamine for 1 day, while Group 5 received a sequential dose of SP for 1 day and AL for 3 days. Sequential administration of AL and SP resulted in a significant (*p* < 0.05) elevation of ALT, AST, ALP, serum total and direct bilirubin levels, urea, creatinine, and HDL. There was a significant (pL0.05) decrease in the serum total protein and albumin. Notably, HDL levels increased significantly in the SP → AL group (*p* < 0.05), while other lipid parameters showed sequence-specific significant changes compared to the control. Sequential administration, particularly the SP → AL sequence, was observed to have more pronounced effects on hepatorenal biomarkers compared to independent administration. These findings are relevant, especially in malaria-endemic regions where unregulated self-medication and drug switching are rampant.

## INTRODUCTION

Malaria is one of the most devastating infectious diseases with an annual global burden of over 200 million, alongside hundreds of thousands of deaths reported each year and apprximately 95% occurring in the WHO African Region [19,40]. It is a protozoan disease transmitted by Anopheles female mosquitoes and results from the infection of a vulnerable host by Plasmodium parasites, of which over 120 species are known, with only five (P. falciparum, P. vivax, P. malariae, P. ovale, P. knowlesi) causing malarial infections in humans. Mortality is predominantly attributed to infection with P. falciparum and P. vivax, heretofore associated with uncomplicated malaria, has recently been documented as a causative agent of severe human infection [49]. In 2024, it is estimated that there were 282 million cases of malaria in 80 endemic countries, an increase of 9 million cases from 2023, and this is determined by various complex factors, including population growth, conflict, climate events, and health system disruptions [50].

The primary impediment to successful malaria treatment is the proliferation of parasite resistance to antimalarial drugs in Africa. Consequently, the World Health Organization (WHO) recommended that antimalarial therapies be deployed as combination therapies (CTs) rather than the usual monotherapies [38]. Coartem®, an artemisinin combination therapy containing Artemether-Lumefantrine (AL), is the first-line drug recommended for the treatment of acute and complicated malaria fever in patients with a minimum body weight of 5 kg, and it is widely used across malaria-endemic regions [26]. Clinical trials showed that Coartem® is effective, safe, and well-tolerated against multidrug-resistant Plasmodium falciparum [39]. Artemisinin derivatives rapidly reduce parasite biomass, achieving up to a 10,000-fold reduction per asexual cycle [29].

Despite the scale-up of antimalarial interventions and the resultant reduction in malaria transmission in some areas, the morbidity and mortality of malaria disease are still high in several areas of sub-Saharan Africa [30]. Artemether–Lumefantrine and Sulfadoxine–Pyrimethamine (SP) are both recommended by the WHO and have contributed to reductions in malaria-related morbidity and mortality in different populations [20,31]. Artemisinin-based combination therapies (ACTs) combine a fast-acting artemisinin derivative with a long-acting drug to prevent recrudescence and drug resistance [38]. The emergence of antimalarial drug resistance has led various countries in sub-Saharan Africa to adopt the World Health Organization (WHO) recommended artemisinin-based combination therapy for treatment of uncomplicated P. falciparum malaria [39]. In routine clinical practice, treatment failure, self-medication, drug switching, and mass drug administration programs may result in the independent or sequential use of different antimalarial agents. However, the biochemical consequences of their sequential administration, particularly the effect of drug order, are still not well understood.

The liver is the most sensitive predictor of chemical-induced toxicity because of its involvement in metabolism, detoxification, and storage of drugs and their metabolites. It is an important target organ for drug-induced injury in mammals. The leakage of cytosolic enzymes such as alanine aminotransferase (ALT), aspartate aminotransferase (AST), and alkaline phosphatase (ALP) is closely related to the distortion of hepatocyte membrane integrity [52]. These enzymes are commonly used as markers of hepatocellular stress. The kidney is affected because of its significant role in drug elimination and maintaining metabolic balance. Changes in serum urea and creatinine levels are used as early markers of kidney function impairment. These modulations can be detected before any histopathological changes in the kidney, making them markers of functional organ stress [17].

Artemisinin derivatives such as artemether are metabolized in the liver by cytochrome P450 enzymes. These drugs have been shown to modulate enzyme activity through induction mechanisms [51]. Modulation of drug metabolism dynamics by sequential exposure to antimalarial drugs may alter drug clearance, leading to the accumulation of reactive metabolites. This can affect drug-metabolizing pathways, leading to different effects in the liver and kidneys depending on the sequence of drug administration [53].

Considering the widespread use of Artemether–Lumefantrine and Sulfadoxine–Pyrimethamine and the likelihood of their sequential administration in malaria-endemic regions, it is therefore pertinent to evaluate the biochemical consequences of such exposure, especially early biochemical indicators of hepatorenal stress. This study is the first Therefore, the objective of this study was to ascertain sequence-specific variations in the biochemical markers of the liver and kidneys after both independent and sequential administrations of Artemether–Lumefantrine and Sulfadoxine–Pyrimethamine in Wistar rats, thereby identifying early indicators of organ stress that may be reversible.

## MATERIALS AND METHODS

### Experimental Animals

Thirty male albino Wistar rats weighing 190-280g were procured from the Department of Zoology Animal House, Faculty of Science, University of Uyo (UNIUYO), Akwa Ibom State. The Animals were weighed, labeled, and kept in cages at the Animal House, Faculty of Basic Medical Sciences (FBMS), UNIUYO, Akwa Ibom State, Nigeria. The rats were acclimatized in an optimum pathogen-free environment and maintained a 12hr light/dark cycle (light on at 6:00 a.m.) at 25-27 °C for 2 weeks prior to the start of the experiment, to allow free and unhindered access to diet and water. The animals were separated into cages with dimensions of 23 cm in length, 10.3 cm in width, and 13 cm in height. The cages were properly maintained by changing the sawdust and leftover feed daily. The rats were fed with rat pellets supplied by Grand Cereals Ltd, Onitsha, Nigeria, and were maintained under standard conditions. Also, the Department of Biochemistry, University of Uyo, Nigeria, Animal Experiment Committee approved the protocol for all animal experiments conducted in this study. All procedures followed the standards established in the NIH Guide for the Care and Use of Laboratory Animals (NIH Publication No. 83-123, revised 1985). The Postgraduate Committee of the Faculty of Basic Medical Sciences at the University of Uyo, Nigeria, approved this study, which received Registration number UU_FBMSREC_2025_002. Efforts were made to reduce the pain caused to the animals and the number of animals required for the study.

### Drugs Acquisition

Artemether-Lumefantrin 80/480mg/Kg (Coartem®, Novartis, Switzerland) and Sulphadoxine-Pyrimethamine 500/25mg/kg (Wapmalar®, Ipca Laboratories, India) were sourced from Somboson Pharmacy, Uyo, Akwa Ibom State, Nigeria.

### Experimental Design

The adult male Wistar rats were weighed, marked, and divided into five groups of six (6) per group. Group 1 served as the control with no drugs administered. Groups 2-5 were administered the following;

| Group | Treatment | Dose (mg/kg<br>bw) | Duration | Dosing Schedule |
| --- | --- | --- | --- | --- |
| 1 | NC – Distilled Water | 5 mL/kg | 4 days | 5 mL/kg/day |
| 2 | Artemether–Lumefantrine<br>(AL) | 8 | 3 days | Day 1–3: 8 mg/kg/day |
| 3 | Sulfadoxine–Pyrimethamine<br>(SP) | 0.079 | 1 day | Day 1: 0.079 mg/kg |
| 4 | AL → SP (Sequential) | AL 8 + SP<br>0.079 | 4 days | Day 1–3: AL 8 mg/kg/day; Day 4:<br>SP 0.079 mg/kg |
| 5 | SP → AL (Sequential) | SP 0.079 + AL<br>8 | 4 days | Day 1: SP 0.079 mg/kg; Day 2–4:<br>AL 8 mg/kg/day |

### Administration of Experimental Drugs

The therapeutic doses of AL and SP were determined and administered orally at 8mg/Kg body weight (bw). The tablets were powdered using a laboratory mortar and pestle. For AL, a stock solution of 5.6mg/mL was made by dissolving 560mg of the powder in 100 mL of water. From this, a further serial dilution was performed to obtain concentrations tolerated by the experimental animals based on their weight. For SP, 3 tablets were dissolved in 60ml of water and further diluted to suit the experimental animals.

### Animal Sacrifice and Preparation of Sera for Analysis

On the fourth day, animals in both experimental and control groups were placed in a glass jar containing cotton wool dipped into ketamine for general anesthesia. The animals were dissected, and blood samples were obtained by cardiac puncture using a sterile needle and syringe and transferred into non–heparinized sample tubes. After coagulation, the blood samples were spun at 2000 rpm for 10 minutes using a bench-top centrifuge (MSE Minor, England). The sera were carefully transferred to clean sample tubes and stored at 40 °C in a refrigerator for further analysis.

### Biochemical Analyses

Serum AST was determined by the Kinetic Method [43], ALT by the colorimetric endpoint method [44], and total and direct bilirubin by Doumas et al. [10]. Creatinine and urea were assayed spectrophotometrically [16,41]. Serum Potassium, Sodium, and Chloride were measured using an automated ion-selective electrode machine (Lanwing LWE60D, Germany), while bicarbonate was determined spectrophotometrically [43]. Serum Lipid Profile, including triglycerides, total cholesterol, and high-density lipoprotein cholesterol, was assayed using the Fortress Assay Kit. Very low-density lipoprotein cholesterol and Low-density lipoprotein cholesterol were calculated using Friedwald’s formula [13].

### Statistical Analyses

Data were analyzed using SPSS v20.0 and expressed as mean ± SEM. Student’s t-test, ANOVA, and LSD post hoc tests were used for group comparisons. Differences were considered statistically significant at (*p* < 0.05).

## RESULTS

### Effect on liver function

Results from liver function analyses revealed that sequential treatment with Artemether-Lumefantrine and Sulfadoxine-pyrimethamine significantly (p 0.05) increased ALT, AST, ALP, and serum total and direct bilirubin levels compared with the control. There were significantly (*p* < 0.05) decreased levels of serum total protein and albumin following sequential treatment with Artemether-Lumefantrine and Sulfadoxine-pyrimethamine compared with the control, as shown in Table 1 and Figure 1.

**Figure 1.**
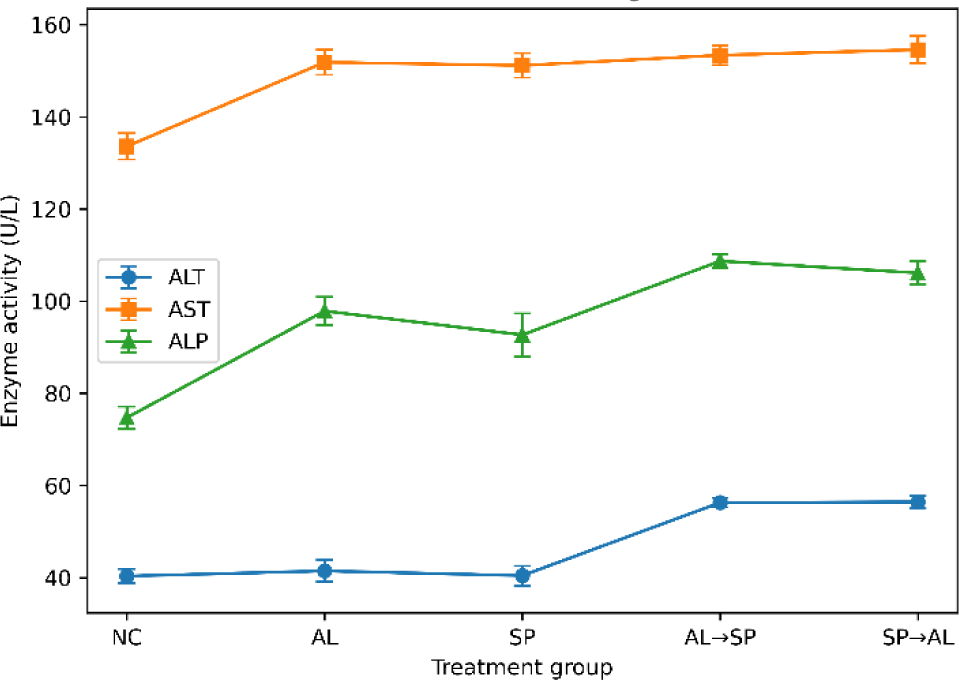
Liver enzyme biomarkers following AL and SP administration

**Figure 2.**
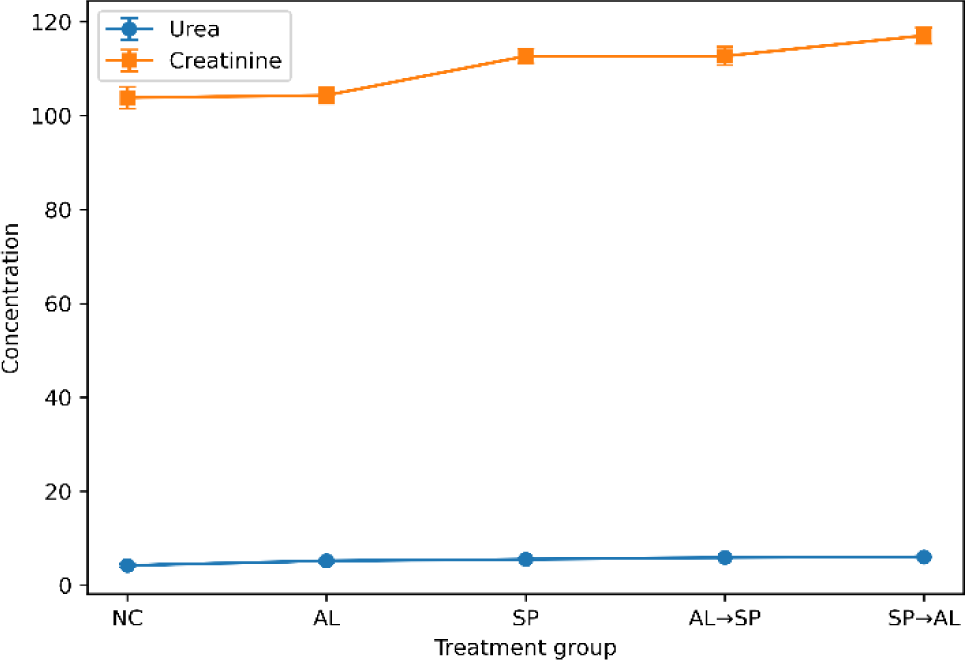
Renal biochemical markers following AL and SP administration

**Figure 3.**
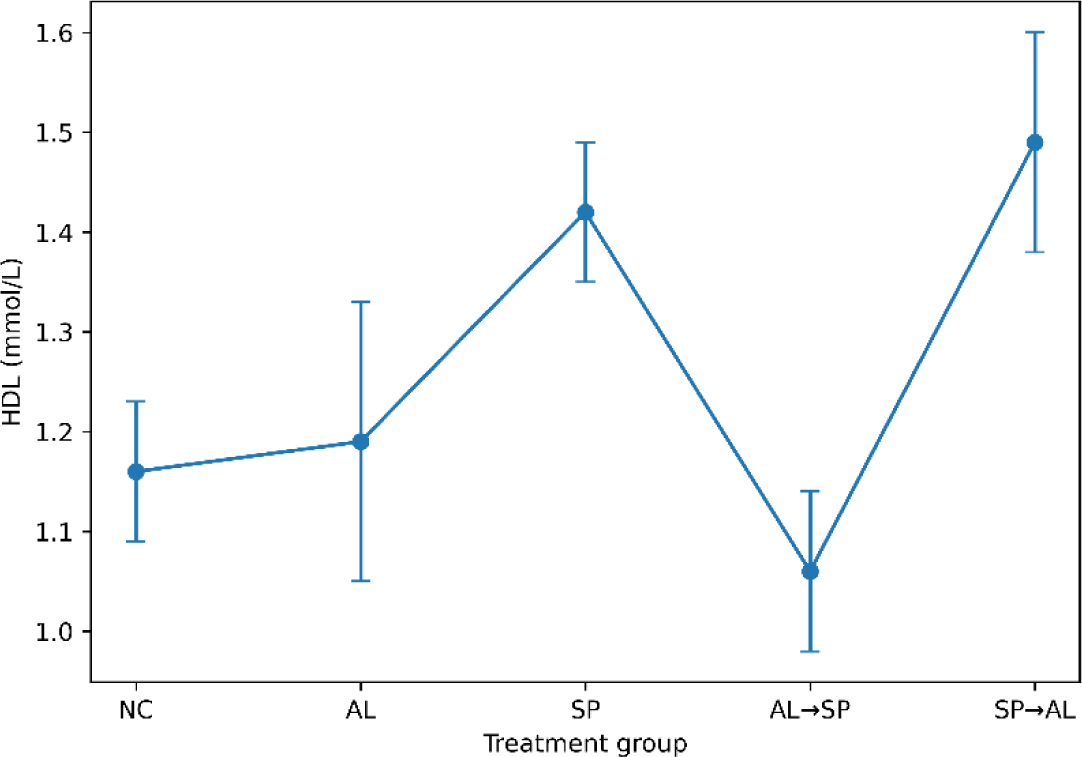
Serum HDL levels following independent and sequential treatme Effects of independent and sequential administration of Artemether–Lumefantrine (AL) and Sulfadoxine–Pyrimethamine (SP) on liver enzymes (ALT, AST, ALP), renal markers (urea, creatinine), and serum high-density lipoprotein (HDL) levels in Wistar rats. Data are expressed as mean ± SEM (n = 6).

**TABLE 1:**
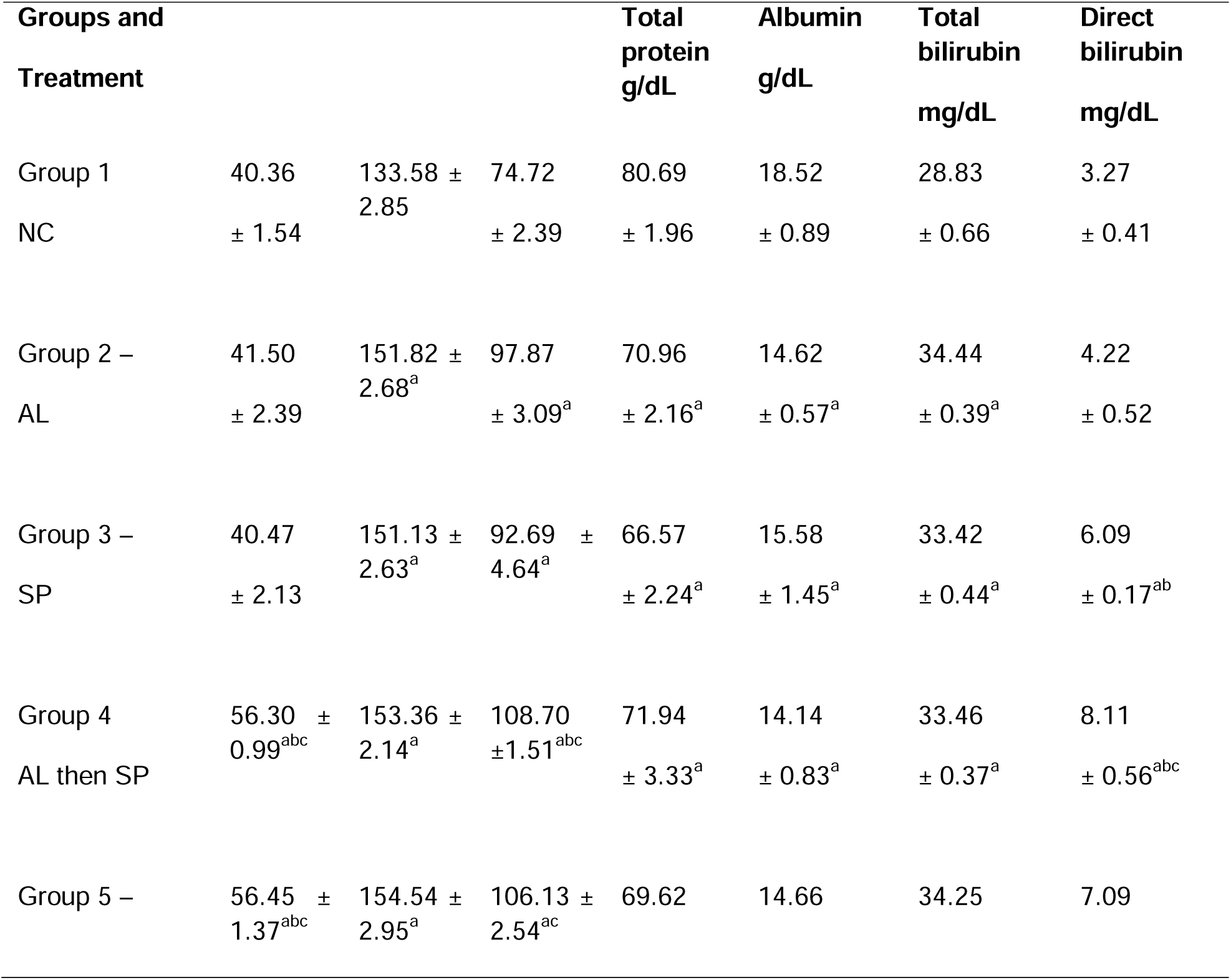
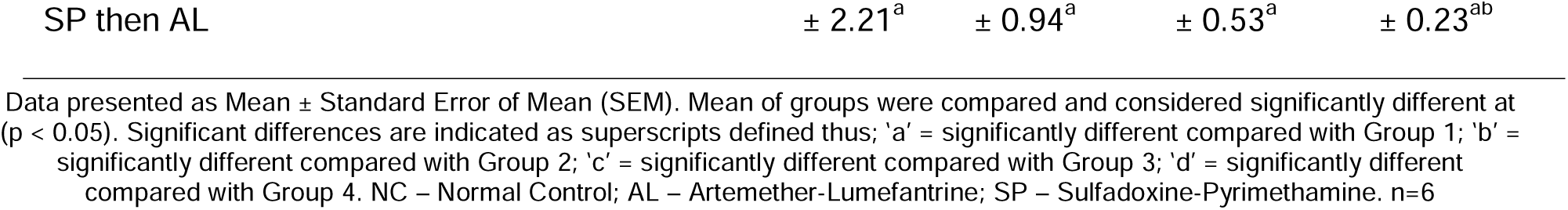
Biomarkers of liver function in Wistar rats sequentially treated with Artemether-Lumefantrine and Sulfadoxine-Pyrimethamine.

| Groups and Treatment |  |  |  | Total protein g/dL | Albumin g/dL | Total bilirubin mg/dL | Direct bilirubin mg/dL |
| --- | --- | --- | --- | --- | --- | --- | --- |
| Group 1 | 40.36 | 133.58 ± 2.85 | 74.72 | 80.69 | 18.52 | 28.83 | 3.27 |
| NC | ± 1.54 |  | ± 2.39 | ± 1.96 | ± 0.89 | ± 0.66 | ± 0.41 |
| Group 2 – | 41.50 | 151.82 ± 2.68<sup>a</sup> | 97.87 | 70.96 | 14.62 | 34.44 | 4.22 |
| AL | ± 2.39 |  | ± 3.09<sup>a</sup> | ± 2.16<sup>a</sup> | ± 0.57<sup>a</sup> | ± 0.39<sup>a</sup> | ± 0.52 |
| Group 3 – | 40.47 | 151.13 ± 2.63<sup>a</sup> | 92.69 ± 4.64<sup>a</sup> | 66.57 | 15.58 | 33.42 | 6.09 |
| SP | ± 2.13 |  |  | ± 2.24<sup>a</sup> | ± 1.45<sup>a</sup> | ± 0.44<sup>a</sup> | ± 0.17<sup>ab</sup> |
| Group 4 | 56.30 ± 0.99<sup>abc</sup> | 153.36 ± 2.14<sup>a</sup> | 108.70 ± 1.51<sup>abc</sup> | 71.94 | 14.14 | 33.46 | 8.11 |
| AL then SP |  |  |  | ± 3.33<sup>a</sup> | ± 0.83<sup>a</sup> | ± 0.37<sup>a</sup> | ± 0.56<sup>abc</sup> |
| Group 5 – | 56.45 ± 1.37<sup>abc</sup> | 154.54 ± 2.95<sup>a</sup> | 106.13 ± 2.54<sup>ac</sup> | 69.62 | 14.66 | 34.25 | 7.09 |

Data presented as Mean ± Standard Error of Mean (SEM). Mean of groups were compared and considered significantly different at ( $p < 0.05$ ). Significant differences are indicated as superscripts defined thus; 'a' = significantly different compared with Group 1; 'b' = significantly different compared with Group 2; 'c' = significantly different compared with Group 3; 'd' = significantly different compared with Group 4. NC – Normal Control; AL – Artemether-Lumefantrine; SP – Sulfadoxine-Pyrimethamine. n=6

### Effects on Kidney Function

Serum concentrations of urea, creatinine, and electrolytes were analyzed to ascertain kidney function. Significant (p 0.05) increases in serum urea and creatinine levels were observed in the sequential treatment with Artemether-Lumefantrine and Sulfadoxine-pyrimethamine compared with the control. There were no significant differences in serum electrolyte levels (sodium, potassium, chloride, and bicarbonate) when all treatment groups were compared with the control, as shown in Table 2 and Figure 2.

**TABLE 2:**
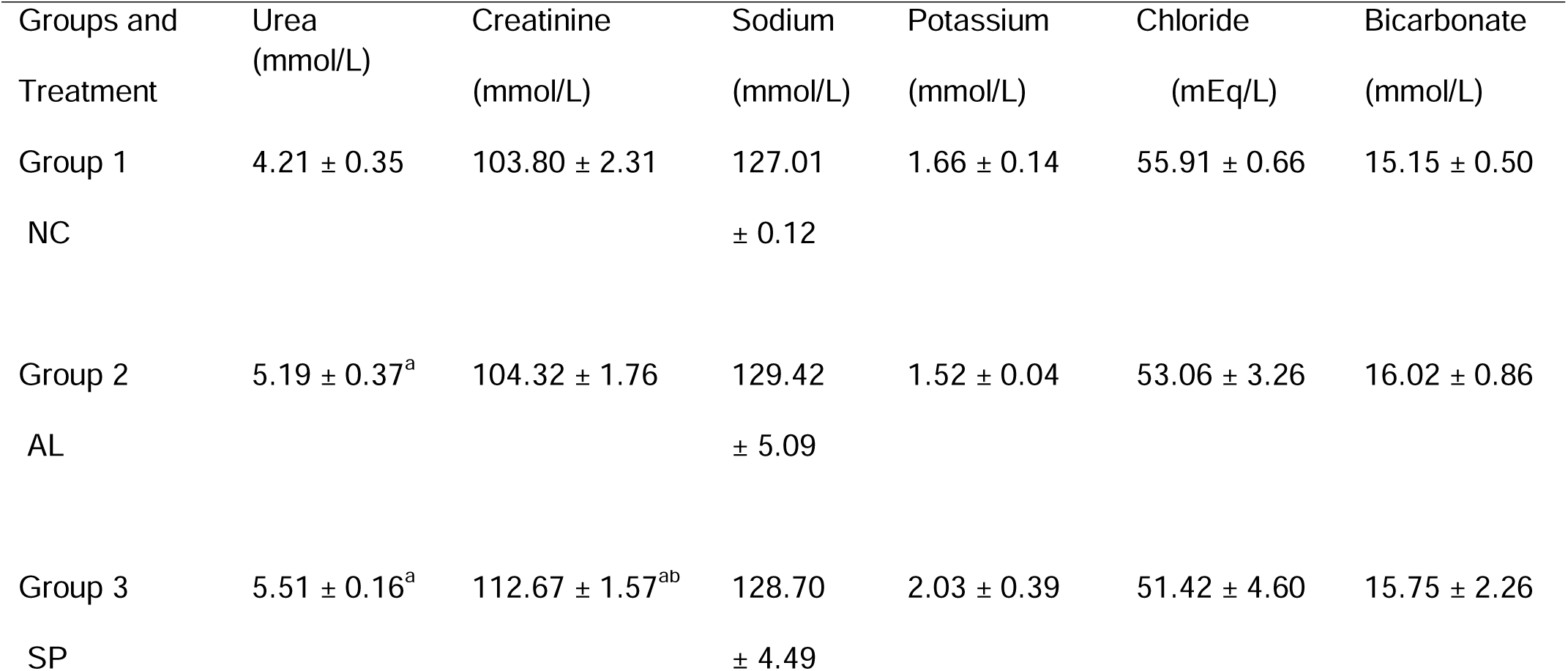
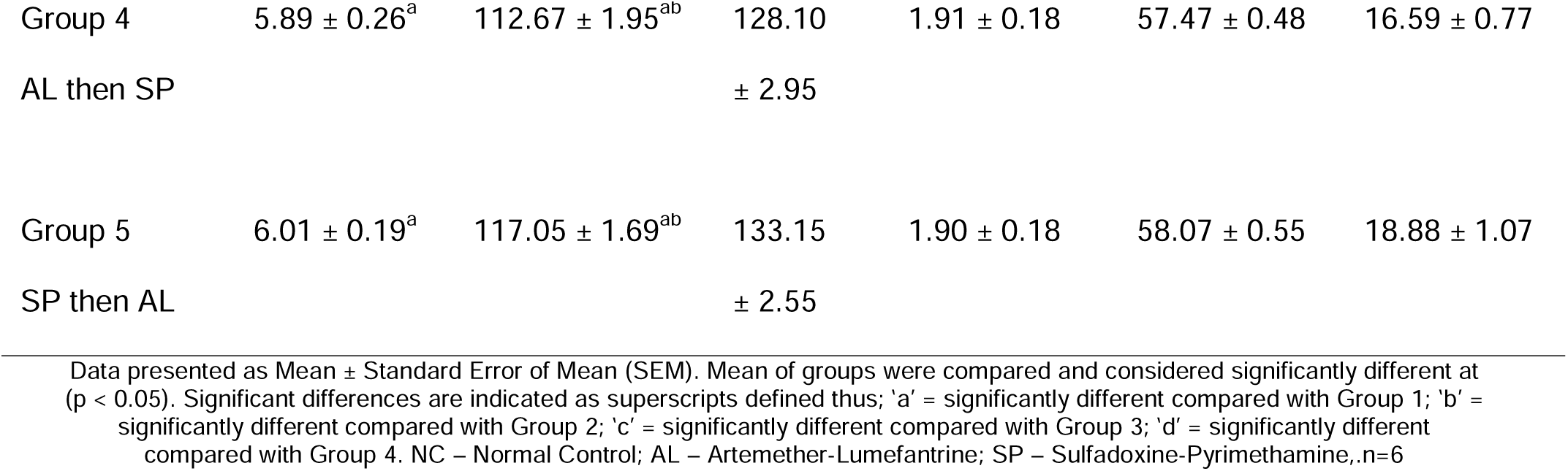
Biomarkers of kidney function in Wistar rats sequentially treated with Artemether-Lumefantrine and Sulfadoxine-Pyrimethamine.

| Groups and Treatment | Urea (mmol/L) | Creatinine (mmol/L) | Sodium (mmol/L) | Potassium (mmol/L) | Chloride (mEq/L) | Bicarbonate (mmol/L) |
| --- | --- | --- | --- | --- | --- | --- |
| Group 1<br>NC | 4.21 ± 0.35 | 103.80 ± 2.31 | 127.01 ± 0.12 | 1.66 ± 0.14 | 55.91 ± 0.66 | 15.15 ± 0.50 |
| Group 2<br>AL | 5.19 ± 0.37<sup>a</sup> | 104.32 ± 1.76 | 129.42 ± 5.09 | 1.52 ± 0.04 | 53.06 ± 3.26 | 16.02 ± 0.86 |
| Group 3<br>SP | 5.51 ± 0.16<sup>a</sup> | 112.67 ± 1.57<sup>ab</sup> | 128.70 ± 4.49 | 2.03 ± 0.39 | 51.42 ± 4.60 | 15.75 ± 2.26 |
| Group 4 | 5.89 ± 0.26<sup>a</sup> | 112.67 ± 1.95<sup>ab</sup> | 128.10 | 1.91 ± 0.18 | 57.47 ± 0.48 | 16.59 ± 0.77 |
| AL then SP |  |  | ± 2.95 |  |  |  |
| Group 5 | 6.01 ± 0.19<sup>a</sup> | 117.05 ± 1.69<sup>ab</sup> | 133.15 | 1.90 ± 0.18 | 58.07 ± 0.55 | 18.88 ± 1.07 |
| SP then AL |  |  | ± 2.55 |  |  |  |

Data presented as Mean ± Standard Error of Mean (SEM). Mean of groups were compared and considered significantly different at ( $p < 0.05$ ). Significant differences are indicated as superscripts defined thus; 'a' = significantly different compared with Group 1; 'b' = significantly different compared with Group 2; 'c' = significantly different compared with Group 3; 'd' = significantly different compared with Group 4. NC – Normal Control; AL – Artemether-Lumefantrine; SP – Sulfadoxine-Pyrimethamine, n=6

### Effect on lipid parameters

The total cholesterol (TCHOL), triacylglyceride (TAG), high-density lipoprotein (HDL), low-density lipoprotein (LDL), and very low-density lipoprotein (VLDL) in Wistar rats sequentially administered Artemether-Lumefantrine and Sulfadoxine-pyrimethamine are presented in Table 3 and Figure 2. Serum high-density cholesterol concentration was significantly (*p* < 0.05) increased with sequential treatment with Sulfadoxine-pyrimethamine and Artemether-Lumefantrine compared with the control. Group 5 (SP then AL) showed significant differences (*p* < 0.05) in the serum TCHOL, TAG, LDL, and VLDL concentrations following sequential treatment with Sulfadoxine-pyrimethamine and Artemether-Lumefantrine when compared to the independent AL treatment group. Additionally, Group 4 (AL then SP) showed a significant difference in Total Cholesterol compared to the independent SP group. Notably, HDL levels were significantly increased (*p* < 0.05) in the SP then AL sequence compared to the control, independent AL, and AL then SP groups.

**TABLE 3:** Lipid profile of Wistar rats sequentially treated with Artemether-Lumefantrine and Sulfadoxine-Pyrimethamine.

| Groups and Treatment | TCHOL<br>(mmol/L) | TAG<br>(mmol/L) | HDL<br>(mmol/L) | LDL<br>(mmol/L) | VLDL<br>(mmol/L) |
| --- | --- | --- | --- | --- | --- |
| Group 1<br>NC | 2.49 ± 0.06 | 1.09 ± 0.07 | 1.16 ± 0.07 | 0.83 ± 0.06 | 0.50 ± 0.03 |
| Group 2<br>AL | 2.91 ± 0.22 | 0.88 ± 0.07<sup>a</sup> | 1.19 ± 0.14 | 1.33 ± 0.11<sup>a</sup> | 0.40 ± 0.03<sup>a</sup> |
| Group 3<br>SP | 3.04 ± 0.22<sup>a</sup> | 1.06 ± 0.07 | 1.42 ± 0.07 | 1.14 ± 0.16 | 0.48 ± 0.03 |
| Group 4<br>AL then SP | 2.43 ± 0.06<sup>bc</sup> | 1.02 ± 0.05 | 1.06 ± 0.08<sup>c</sup> | 0.91 ± 0.08<sup>b</sup> | 0.46 ± 0.02 |
| Group 5<br>SP then AL | 2.94 ± 0.11<sup>d</sup> | 1.15 ± 0.08<sup>b</sup> | 1.49 ± 0.11<sup>abd</sup> | 0.93 ± 0.11<sup>b</sup> | 0.52 ± 0.03<sup>b</sup> |

Data presented as Mean ± Standard Error of Mean (SEM). Mean of groups were compared and considered significantly different at ( $p < 0.05$ ). Significant differences are indicated as superscripts defined thus; 'a' = significantly different compared with Group 1; 'b' = significantly different compared with Group 2; 'c' = significantly different compared with Group 3; 'd' = significantly different compared with Group 4. NC – Normal Control; AL – Artemether-Lumefantrine; SP – Sulfadoxine-Pyrimethamine; TCHOL= Total Cholesterol, HDL = High Density Lipoprotein, TAG = Triacylglyceride, VLDL = Very Low Density Lipoprotein, LDL = Low Density

Effects of independent and sequential administration of Artemether–Lumefantrine (AL) and Sulfadoxine–Pyrimethamine (SP) on liver enzymes (ALT, AST, ALP), renal markers (urea, creatinine), and serum high-density lipoprotein (HDL) levels in Wistar rats. Data are expressed as mean ± SEM (n = 6).

## DISCUSSION

Hepatocyte membrane distortion is associated with increased permeability and subsequent leakage of intracellular enzymes into the circulation, as evidenced by elevations in serum alanine aminotransferase (ALT), aspartate aminotransferase (AST), and alkaline phosphatase (ALP), which are generally considered biomarkers of hepatocellular injury [37]. Among these, ALT is usually the most reliable indicator of hepatocellular damage, whereas AST, although found in hepatocytes, is also present in extrahepatic tissues such as cardiac and skeletal muscles, kidneys, and testes, making its elevation not only hepatic in origin [33].

In this study, AST levels were elevated in all treatment groups and were statistically significant compared with the normal control, indicating systemic enzyme leakage following both independent and sequential drug administration. Similarly, ALP levels were significantly higher in treated groups, especially sequentially treated animals. However, ALT activity was significantly elevated following sequential administration (Groups 4 and 5) but not with independent administration of AL or SP (Groups 2 and 3), suggesting that combined exposure exerts greater hepatocellular toxicity than single-dose therapy. This is corroborated by other studies showing that combined malaria regimens resulted in elevated ALT levels [22,45,46,47]. Meanwhile, in a Preclinical study in rodent malaria models conducted by Okafor et al. [48], it was observed that artemisinin-based combination therapies induce histopathological changes in multiple organs, including the liver and kidneys.

Furthermore, a modest increase in total and direct bilirubin was observed, mainly in the sequential treatment groups, indicating some impairment of hepatobiliary processing and conjugation. Although this may reflect early functional disturbance rather than fixed liver damage. Interestingly, single SP administration caused relatively less enzyme induction than AL, which may explain why SP continues to be safely used in intermittent malaria prevention programs.

The sequence-specific biochemical effects could be partly explained by the liver’s metabolism of these drugs. Artemether, for example, is extensively metabolized in the liver by CYP3A4 and CYP2B6. Some studies suggest that artemisinin induces these enzymes through auto-induction, which could alter the metabolism and clearance of a drug administered afterward [42,6]. SP, by contrast, has slower clearance and longer systemic exposure. Drugs with extended elimination phases are thought to be more prone to enzyme-mediated interactions, potentially leading to metabolic competition and accumulation of reactive intermediates [34,8]. This may have contributed to the more pronounced effect on group 5. Conversely, the administration of AL before SP may have induced the metabolism of the second drug so rapidly that the biochemistry was altered to a greater extent. Once again, this is speculative, but it is consistent with the results of previous studies on the pharmacokinetics-toxicodynamics of drug interactions [6, 42]. Sequential administration also seemed to affect renal biomarkers. Serum urea and creatinine were significantly elevated in the SP → AL group, suggesting reduced glomerular filtration and possible nephrotoxicity [11,15]. Electrolytes remained generally within normal limits, indicating that the kidney’s regulatory mechanisms remained functional. The fluctuations in electrolyte levels could be due to a slight decrease in kidney efficiency, as observed in earlier studies of antimalarial administration [12,14,32].

Additionally, in this study, there were sequence-dependent changes in lipid parameters. HDL increased significantly in the SP → AL group compared to the control, independent AL, and AL → SP groups. Interestingly, independent administration of SP (Group 3) resulted in significantly elevated Total Cholesterol compared to the control. In contrast, the sequential AL → SP group (Group 4) showed significantly lower Total Cholesterol levels than both independent AL and SP treatments. Furthermore, while independent AL treatment (Group 2) caused a significant reduction in TAG and VLDL compared to the control, sequential SP → AL administration resulted in significantly higher levels of these lipids compared to the AL group. This may be a risk factor for atherosclerosis or coronary heart disease [25,12,28,5,21]. This suggests that the order of drug administration could serve as an early, sensitive indicator of organ stress in Wistar rats and hints that sequence effects may require careful consideration in real-world settings, particularly where drug switching or self-medication is common, to optimize the hepatorenal safety of malaria chemotherapy.

## Conclusion

This study shows that the biochemical effects of Artemether–Lumefantrine and Sulfadoxine–Pyrimethamine are influenced by the sequence of administration in male Wistar rats. Independent administration of Artemether–Lumefantrine (once daily for three days) or a single dose of Sulfadoxine–Pyrimethamine produced only mild and largely physiological changes in serum biochemical indices. However, sequential exposure resulted in more pronounced alterations, with the Sulfadoxine–Pyrimethamine-Artemether–Lumefantrine sequence showing higher elevations in hepatic and renal biochemical biomarkers. This study, therefore, provides novel evidence that should inform safety protocols and pharmacovigilance for antimalarial therapy. It is worth noting that a limitation of this study is the absence of a histopathological assessment to corroborate our findings.

## Ethical Approval

This study was approved by the Postgraduate Ethical Committee of University of Uyo, Uyo, Nigeria, with registration number: UU_FBMSREC_2025_002.

## Informed Consent

This is not applicable to this article as no human subjects were involved in this study.

## References

1. Abdelghffar EA. Pathophysiological effects of Tamiflu on liver and kidneys of male rats. Beni-Suef Univ J Basic Appl Sci. 2022;11(1):1–14.

2. Abolaji AO, Eteng MU, Omonua O, Adenrele Y. Influence of co-administration of artemether and lumefantrine on selected plasma biochemical and erythrocyte oxidative stress indices in female Wistar rats. Hum Exp Toxicol. 2013;32(2):206–15.

3. Al-Faris OJ, Al-Shawi NN, Kako MD. Possible cardiac adverse effects induced by therapeutic doses of ciprofloxacin in juvenile rats. Iraqi J Pharm Sci. 2012;21(2):94–7.

4. Amorha KC, Ugwuowo OB, Ayogu EE, Nduka SO, Okonta MJ. Evaluation of the hepatic effect of concomitant administration of ciprofloxacin and some antimalarial drugs in Plasmodium berghei–infected mice: An in vivo study. Pak J Pharm Sci. 2018;31(5):1805–11.

5. Amsterdam EA, Wenger NK, Brindis RG, Casey DE, Ganiats TG, Holmes C, et al. AHA/ACC guideline for the management of patients with non-ST-elevation acute coronary syndromes: A report of the American College of Cardiology/American Heart Association Task Force on Practice Guidelines. J Am Coll Cardiol. 2014;64(24):2654–87.

6. Asimus S, Elsherbiny D, Hai TN, Jansson B, Huong NV, Petzold MG, et al. Artemisinin antimalarials moderately affect cytochrome P450 enzyme activity in healthy subjects. Fundam Clin Pharmacol. 2007;21(3):307–16.

7. Badawy FA, Ali AA, Esmail NS, Helmy AH. Hyperchloremia in critically ill patients in ICU: Review article. Egypt J Hosp Med. 2022;86(1):532–7.

8. Brunton LL, Hilal-Dandan R, Knollmann BC, editors. Goodman & Gilman’s: The Pharmacological Basis of Therapeutics. 13th ed. New York: McGraw-Hill Education; 2018.

9. Chen S, Chiaramonte R. In creatinine kinetics, the glomerular filtration rate always moves the serum creatinine in the opposite direction. Physiol Rep. 2021;9(16): e14957.

10. Doumas BT, Perry BW, Sasse EA, Straumfjord JV. Standardization in bilirubin assays: Evaluation of selected methods and stability of bilirubin solutions. Clin Chem. 1973;19(9):984–993.

11. Edagha IA, Ekpo AJ, Edagha EI, Bassey JV, Nyong TP, Akpan AS, et al. Investigating the comparative effects of six artemisinin-based combination therapies on Plasmodium-induced hepatorenal toxicity. Niger Med J. 2019;60(4):211–8.

12. Etim O, Ekaidem I, Akpan E, Usoh I, Akpan H. Changes in lipid profile and some liver enzymes in rats co-administered with artemether-lumefantrine and ciprofloxacin. Issues Biol Sci Pharm Res. 2018;6(1):1–7.

13. Friedewald WT, Levy RI, Fredrickson DS. Estimation of the concentration of low-density lipoprotein cholesterol in plasma, without use of the preparative ultracentrifuge. Clin Chem. 1972;18(6):499–502.

14. Gallafassi EA, Bezerra MB, Rebouças NA. Control of sodium and potassium homeostasis by renal distal convoluted tubules. Braz J Med Biol Res. 2023;56: e12392.

15. Gowda S, Desai PB, Kulkarni SS, Hull VV, Math AAK, Vernekar SN. Markers of renal function tests. North Am J Med Sci. 2010;2(4):170–3.

16. Henry RJ. Clinical Chemistry: Principles and Techniques. 2nd ed. Hagerstown (MD): Harper & Row; 1974.

17. Idowu ET, Alimba CG, Olowu EA, Otubanjo AO. Artemether lumefantrine treatment combined with albendazole and ivermectin induced genotoxicity and hepatotoxicity through oxidative stress in Wistar rats. Egypt J Basic Appl Sci. 2015; 2:110–9.

18. Lala V, Zubair M, Minter DA. Liver Function Tests. [Updated 2023 Jul 30]. In: StatPearls [Internet]. Treasure Island (FL): StatPearls Publishing; 2025 Jan-. Available from: https://www.ncbi.nlm.nih.gov/books/NBK482489/

19. Lamesgen A, Engidaw M, Gedif G, Gete M, Belay YA. The economic burden of malaria in Africa: A systematic review of cost of illness studies. Malar J. 2025; 24:223.

20. Makanga M, Krudsood S. The clinical efficacy of artemether/lumefantrine (Coartem®). Malar J. 2009;8(Suppl 1): S5.

21. Marz W, Kleber ME, Scharnagl H. HDL-cholesterol: Reappraisal of its clinical relevance. Clin Res Cardiol. 2017;106(9):663–75.

22. Moosavy SH, Eftekhar E, Davoodian P, Ghojazadeh M, Malekzadeh R. AST/ALT ratio, APRI, and FIB 4 compared to FibroScan for the assessment of liver fibrosis in patients with chronic hepatitis B in Bandar Abbas, Hormozgan, Iran. BMC Gastroenterol. 2023; 23:145.

23. Nagami GT. Hyperchloremia – Why and how. Nefrología (Engl Ed). 2016;36(4):347–53.

24. National Kidney Foundation. Clinical practice guideline for the evaluation and management of chronic kidney disease. Kidney Disease Outcomes Quality Initiative (KDOQI) [Internet]. 2024 [cited 2026 Feb 14]. Available from: https://www.kidney.org/sites/default/files/2024-08/ckd_evaluation_classification_stratification.pdf

25. Ndem JI, Sylvanus PU, Bassey UE, Effiong BO, Ewere EG. Assessing the effect of concomitant administration of artemether-lumefantrine and ciprofloxacin on some cardiac parameters in Wistar rats: “The remedial role of vitamin E.” GSC Biol Pharm Sci. 2021;17(01):094–104.

26. Ngasala BE, Mubi M, Warsame M, et al. Efficacy and effectiveness of artemether–lumefantrine after initial and repeated treatment in children < 5 years with acute uncomplicated Plasmodium falciparum malaria in rural Tanzania: A randomized trial. Clin Infect Dis. 2011; 52:873–82.

27. Nkereuwen E, Paul OO, Elias A. Effect of Artemether treatment on plasma lipid profile in malaria. Pharmacol Pharm. 2014;5:646–56.

28. Nna VU, Ofemi OE, Archibong AN, Bassey SC. Alteration in serum lipid profile following separate administration of antimalarial drugs (coartem and chloroquine): A comparative study. Der Pharma Chemica. 2014;6(4):415–21.

29. Nosten F, White NJ. Artemisinin based combination treatment of Plasmodium falciparum malaria. Am J Trop Med Hyg. 2007;77(6 Suppl):181–92.

30. Nwaiwu O, Okorie PN, Orji AU. Persistent malaria transmission in sub-Saharan Africa despite intervention efforts: A systematic review. Infect Dis Poverty. 2023; 12:46.

31. Pécoul B, Chirac P, Trouiller P, Pinel J. Access to essential drugs in developing countries: A lost battle? JAMA. 2008;281(4):361–7.

32. Rodan AR. Regulation of distal nephron transport by intracellular chloride and potassium. Nephron. 2023;147(3–4):203–11.

33. Shipman AR, Shipman KE. Investigative algorithms for disorders affecting plasma transaminases (aspartate transaminase and alanine transaminase)—A narrative review. J Lab Precis Med. 2024; 9:14.

34. Miyake T, Tsutsui H. Quantitative prediction of human pharmacokinetic drug–drug interactions and drug clearance using humanized liver chimeric mice: A review. Drug Metab Pharmacokinet. 2026; Article 101517.

35. Ugian EA, Dasofunjo K, Nwangwa JN, Asuk AA, Akam MS, Ajing EN, et al. Effect of Artemisinin-Based Combination Therapy on some selected liver function indices of pregnant Wistar albino rats. J Appl Pharm Sci. 2013;3(09):152–4.

36. Ukpanukpong RU, Eteng MU, Dasofunjo K. Antioxidant interactions of pefloxacin, garlic, vitamins C and E on lipid profile level of albino Wistar rats. J Appl Pharm Sci. 2013;3(3):167–70.

37. Umezulike AJ, Maduka SO, Njoku-Oji NN, Okonudo PO, Nwaefulu K, Eluemunor M, et al. Comparative effects of some common vegetable oils on lipid profile and liver function in male Wistar rats. Int J Innov Sci Res Technol. 2021;6(12):1201–12.

38. World Health Organization. Guidelines for the treatment of malaria. 2nd ed. Geneva: WHO; 2010.

39. World Health Organization. Guidelines for malaria. 3rd ed. Geneva: WHO; 2021.

40. World Health Organization. World malaria report 2024: Addressing inequity in the global malaria response. Geneva: WHO; 2024.

41. Wybenga DR, Di Giorgio J, Pileggi VJ. Manual and automated methods for urea nitrogen measurement in whole serum. Clin Chem. 1971;17(10):891–5.

42. Xing J, Kirby BJ, Whittington D, Wan Y, Goodlett DR. Evaluation of P450 inhibition and induction by artemisinin antimalarials in human liver microsomes and primary human hepatocytes. Drug Metab Dispos. 2012;40(9):1757–64.

43. Young DS. Effects of Drugs on Clinical Laboratory Tests. 3rd ed. Washington (DC): AACC Press; 1990.

44. Young DS, Pestaner LC, Gibberman V. Effects of drugs on clinical laboratory tests. Clin Chem. 1975;21(5):1D–432D.

45. Akam U, Okon E, Edem E. Effect of chloroquine, amodiaquine, quinine and halofantrine on serum enzymes. J Pharm Biol Sci. 2013;15(8):33–35.

46. Farombi A, Ademola B, Olatunde M. Effect of coartem and p-alaxin on pregnant albino Wistar rats. Asian Journal of Biochemical Science. 2000;16(7):66–69.

47. Olugbenga MA, Obot O-O, Oluwatoyin HA. Effect of artemisinin-based combination therapy on the liver and kidney of patients attending University Health Centre. Niger J Pharm Appl Sci Res. 2020;7(2):28–32.

48. Okafor UE, Ufele AN, Nwankwo OD. Effects of artemisinin-based combination therapy on histopathology of the liver, kidney and spleen of mice infected with *Plasmodium berghei*. Anim Res Int. 2019;16(3):3519–3528.

49. Matlani M, Kojom LP, Mishra N, et al. Severe *Plasmodium vivax* malaria trends in the last two years: a study from a tertiary care centre, Delhi, India. Ann Clin Microbiol Antimicrob. 2020; 19:49. doi:10.1186/s12941-020-00393-9.

50. World Health Organization. World malaria report 2025: addressing the threat of antimalarial drug resistance. Geneva: World Health Organization; 2025. Available from: https://www.who.int/teams/global-malaria-programme/reports/world-malaria-report-2025

51. Xiong Y, Huang J. Anti-malarial drug: the emerging role of artemisinin and its derivatives in liver disease treatment. Chin Med. 2021 Aug 18;16(1):80. doi: 10.1186/s13020-021-00489-0. PMID: 34407830; PMCID: PMC8371597.

52. Lopez J, Carl A. Burtis and David E. Bruns: Tietz Fundamentals of Clinical Chemistry and Molecular Diagnostics, 7th ed. Elsevier, Amsterdam, 1075 pp, ISBN 978-1-4557-4165-6. Indian J Clin Biochem. 2015 Apr;30(2):243. doi:10.1007/s12291-014-0474-9. PMCID: PMC4393386.

53. Zang M, Zhu F, Li X, Yang A, Xing J. Auto-induction of phase I and phase II metabolism of artemisinin in healthy Chinese subjects after oral administration of a new artemisinin-piperaquine fixed combination. Malar J. 2014 Jun 3;13:214. doi: 10.1186/1475-2875-13-214. PMID: 24889062; PMCID: PMC4055232.

